# A Targeted Reference Database for Improved Analysis of Environmental 16S rRNA Oxford Nanopore Sequencing Data

**DOI:** 10.1101/2024.10.03.616456

**Authors:** Melcy Philip, Tonje Nilsen, Sanna Majaneva, Ragnhild Pettersen, Morten Stokkan, Jessica Louise Ray, Nigel Keeley, Knut Rudi, Lars-Gustav Snipen

## Abstract

The Oxford Nanopore Technologies (ONT) sequencing platform is compact and efficient, making it suitable for rapid biodiversity assessments in remote areas. Despite its long reads, ONT has a higher error rate compared to other platforms, necessitating high-quality reference databases for accurate taxonomic assignments. However, the absence of targeted databases for underexplored habitats, such as the seafloor, limits ONT’s broader applicability for exploratory analysis.

To address this, we propose an approach for building environmentally-targeted databases to improve 16S rRNA gene (16S) analysis using Oxford Nanopore Technologies (ONT), using seafloor sediment samples from the Norwegian coast as an example. We started by using Illumina short-read data to create a database of full-length or near full-length 16S sequences from seafloor samples. Initially, amplicons are mapped to the SILVA database, with matches added to our database. Unmatched amplicons are reconstructed using METASEED and Barrnap methodologies with amplicon and metagenome data. Finally, if the previous strategies did not succeed, we included the short-read sequences in the database. This resulted in AQUAeD-DB, which contains 14 545 16S sequences clustered at 95% identity. Comparative database analysis reveal that AQUAeD-DB provides consistent results for both Illumina and Nanopore read assignments (median correlation coefficient: 0.50), whereas a standard database showed a substantially weaker correlation. These findings also emphasize its potential to recognize both high and low-abundance taxa, which could be key indicators in environmental studies. This work highlights the necessity of targeted databases for environmental analysis, especially for ONT-based studies, and lays foundations for future extension of the database.

## Introduction

The Earth’s surface is primarily covered by oceans, yet we know little about the seafloor, or its microbiota (Carrasco De La Cruz, 2021; Thurber et al., 2014). Microorganisms play a key role in biogeochemical cycling in the global ocean thus the potential importance of microbiota to healthy ecosystem function cannot be overstated (Madsen, 2011). Many aspects of our understanding of soft-sediment microbiota is based on short read amplicon sequencing of the 16S rRNA gene V3 and V4 regions (Ghate et al., 2021; Hoshino et al., 2020; Yang et al., 2022). Advancements in long read sequencing, including Oxford Nanopore Technology (ONT), have enabled rapid analysis of environmental microbiota (Zorz et al., 2023). However, the relatively high error rate of ONT sequencing technology remains a persistent challenge, rendering ONT strategies dependent on reference databases (Amarasinghe et al., 2020). For effective and robust use of ONT technology that generates reads spanning nearly the entire 16S gene, databases of nearly full-length 16S rRNA gene sequences with comprehensive representation of habitat-relevant taxa are needed.

The most widely used and recognized 16S databases are the curated Greengenes and SILVA databases, as well as the non-curated NCBI 16S rRNA database (Federhen, 2011; McDonald et al., 2024; Yilmaz et al., 2013). However, these databases cover only a limited range of environment-associated or marine-related microorganisms, despite housing a large number of reference sequences. Moreover, recent studies have revealed other underlying issues such as taxonomic gaps, partial annotation and redundancy (Cabezas et al., 2024). For instance, the 16S component of the GenBank database consists of 43% chimeric sequences, the presence of which may confound accurate taxonomic assignments (DeSantis et al., 2006; Frenkel-Morgenstern et al., 2012). Furthermore, despite the use of quality control tools, public 16S reference databases still contain 0.2% to 2.5% misidentified species, leading to taxonomic misassignments (Kozlov et al., 2016). In addition, there is a major bias towards characterization of human-associated and highly abundant bacteria, while most of environmental microbial diversity remains uncharacterized (Yang & Iwasaki, 2014). In this context, we fail to notice potentially low-abundant microorganisms, such as nitrifying bacteria, anammox bacteria and methanotrophs, that may be crucial for reflecting the health of the environment (Marlow et al., 2014; Patil et al., 2021; Wu et al., 2019). Benthic microbes, in particular, are underrepresented in databases and our understanding of them is constrained by very few studies (Tatsuhiko Hoshinoa, 2020). The current scenario thus emphasizes the need for a specialized database for seafloor microorganisms, since environmental microbial communities are often missing from current databases.

Studies reveal that employing targeted databases can improve the accuracy and coverage of amplicon-based microbial profiling (Dueholm et al., 2022; Meola et al., 2019; Overgaard et al., 2022). Although ONT sequencing technology is advancing, there are only few bioinformatics tools for ONT 16S profiling (Ciuffreda et al., 2021; Santos et al., 2020; Sissel et al., 2015) apart from the workflow offered by Oxford Nanopore. There are several ongoing efforts to improve taxonomic assignments within ONT-driven studies, with the EMU tool showing promise as an extensively used approach (Curry et al., 2022; Rodríguez-Pérez et al., 2020; Zorz et al., 2023). However, all these tools, require a good reference database for improved taxonomic assignment. This underlines the necessity of an environment-specific or specialized reference database and the associated potential benefits. (F. Escapa et al., 2020; Henderson et al., 2019; Nierychlo et al., 2020; Ritari et al., 2015). Studies have focused on achieving high resolution using Nanopore, given its longer reads covering most of the 16S gene, whereas our study aims to capture the broadest possible diversity from environmental samples (Benítez-Páez et al., 2016; Lemoinne et al., 2023).

The aim of this work is to present an approach for developing a targeted 16S database specifically designed for seafloor sediment or similar environments. This database facilitates microbial profiling and ecosystem status prediction using full-length 16S (V2-V9) ONT sequencing. We have called this database “AQUAeD-DB” in acknowledgment of the research project “AQUAeD” which provided environmental samples for sequencing analysis (see Acknowledgements). Our strategy (Fig. 1) makes use of short-read amplicon data in combination with existing databases, as well as whole genome shotgun sequencing in combination with our recently developed method METASEED (Philip et al., 2024). We also assess and present the effectiveness of this targeted database in comparison to the existing tools and databases for taxonomic assignment of 16S ONT amplicons. As an example, we demonstrate this methodology using sediment samples from the Norwegian coast, with plans to apply it in the monitoring of seafloor sediments to assess the impacts of aquaculture sites across Norway. Additionally, this approach is adaptable for broader and global studies in the future.

**Fig. 1.**
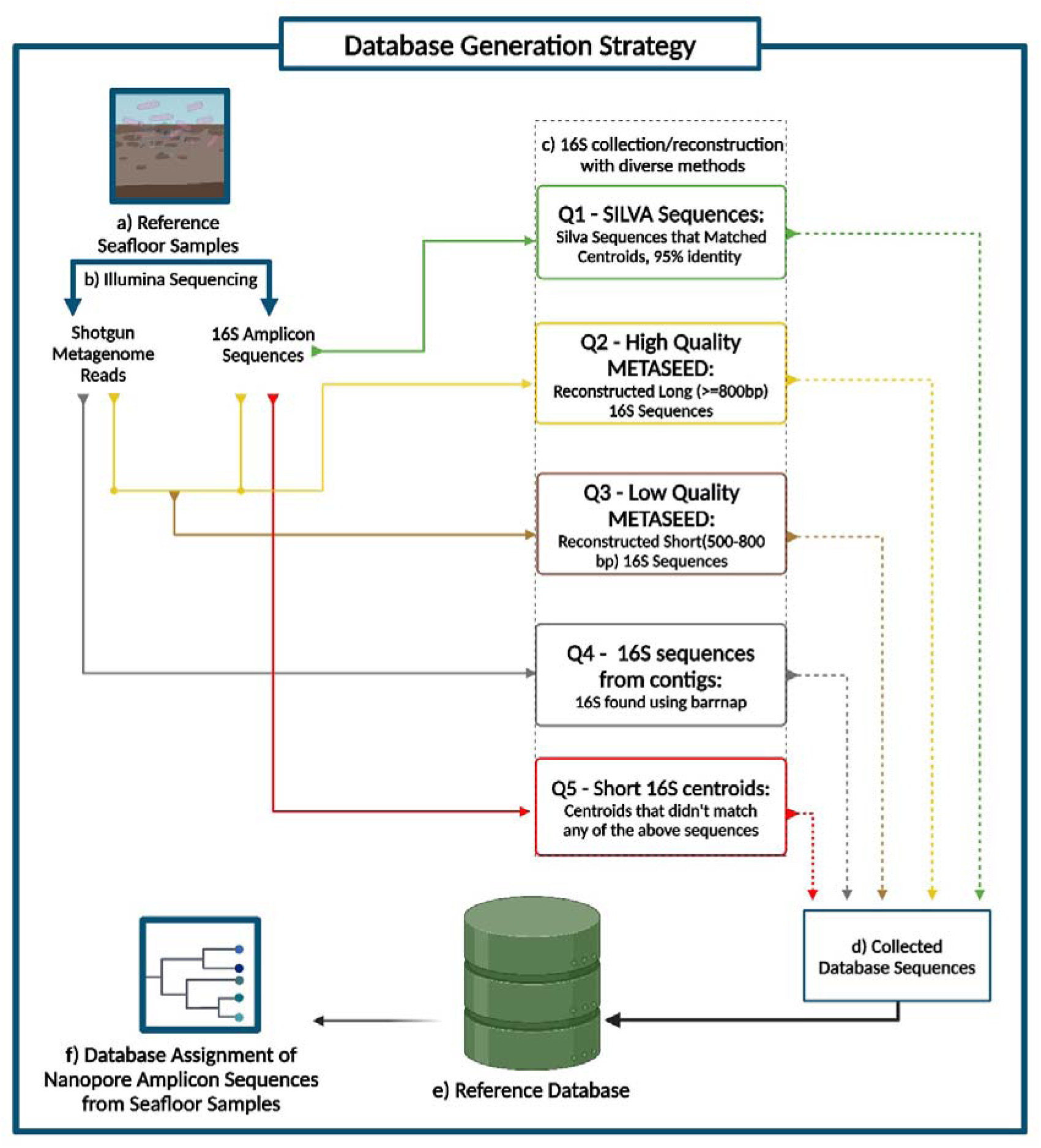
The16S database generation. a) The seafloor samples used in this study were collected from various sites along the Norwegian coast. b) The samples for database generation were sequenced using Illumina, providing shotgun and 16S rRNA amplicon reads, c) Various methods are used to collect or rebuild longer 16S rRNA sequences, which are then graded into different quality levels, from Q1 (best) to Q5 (poorest). These sequences are added to our reference database e) and used to for taxonomic assignment of ONT sequences amplified from geographically proximal seafloor sediment material.

## Materials and Methods

### Data

In this study, we used total of 182 marine sediment samples collected along a north-south gradient along the Norwegian coast as mentioned previously (Nilsen et al., 2024; Pettersen et al., 2022). The sample information and collection details can be found in this study (Nilsen et al., 2024). The data was randomly divided into two subsets: subset A consisted of 94 samples and was used for the database construction; subset B consisted of 88 samples and was used for database comparison.

Subset A was subject to Illumina sequencing, both 16S amplicon and whole genome (metagenome) sequencing. Sediment sample processing was carried out as described in (Pettersen et al., 2022). In brief, sediment DNA was extracted on a KingFisher Flex automated liquid handling platform using the MagAttract PowerSoil DNA KF Kit (Qiagen). The amplification of the V3-V4 region of the 16S rRNA gene was done using primers 341F and 806R (Yu et al., 2005). Individual amplicon PCR products were dual-indexed using different combinations of Illumina forward and reverse primers with unique barcode tags. The library pools were sequenced a MiSeq v3 platform at the Norwegian Sequencing Centre in Oslo. Shotgun sequencing was performed on an Illumina Novaseq 6000 platform at Novogene, UK facility using the protocol stated in (Gourlé et al., 2018).

Subset B samples were subjected to 16S amplicon sequencing using both Illumina and ONT sequencing platforms. Library preparation for Illumina sequencing of Subset B samples was performed as described above. ONT sequencing libraries were prepared using the Oxford Nanopore Sequencing kit (SQK-LSK114-XL). The first amplification was performed using modified 16S forward and reverse primers (Genetic Analysis, Norway) with the following PCR program: 95°C for 15 min, 30 cycles of 95°C for 30 sec, 55°C for 30 sec and 72°C for 80 sec. Amplicons were purified with 1:1 Ampure XP paramagnetic beads (Beckman Coulter) before indexing PCR. Between 10 and 60 ng of purified full-length 16S amplicons was added to the PCR mixture, in addition to 0.2 µM of a unique PCR barcode (ONT cat.nr. EXP-PBC096). The barcoding amplification was done using the following program: 95°C for 15 min, 12 cycles of 95°C for 30 sec, 62°C for 15 min and 65°C for 2 min, followed by 65°C for 10 min. Barcoded PCR products were concentration-normalized and pooled in equimolar ratios, the library from which was purified with 1:1 Ampure XP beads before sequencing on a MinION flow cell. The samples were sequenced at Eurofins Genomics, Germany.

### Amplicon data processing

The 16S amplicon sequencing data from subset A were processed with VSEARCH (Rognes et al., 2016) for clustering into Operational Taxonomic Units (OTUs) using a 95% identity threshold. ONT reads are longer than Illumina reads but still have error levels making it difficult to go much above a 95% identity resolution for the 16S sequences. Because the subset Illumina sequence data will be used to generate reference sequences for error-prone ONT sequences in subset B, a lower 95% resolution was chosen as opposed to a higher value or a denoising approach. In addition, a recent study of very high diversity environments like seafloor sediments demonstrated unstable compositional profiles when using high-resolution denoising of illumina reads (Nilsen et al., 2024).

The Illumina paired-end reads from subset B were merged and de-replicated by VSEARCH, but not clustered. Only reads with at least five copies per sample were used, since these are most likely without sequencing error. Nanopore reads from the same subset B samples were filtered based on their length, which ranged between 800 – 1 400 bp.

### SILVA alignments

The OTU represents the sequences of around 420 - 440 bases, which corresponds to the lengths of the amplicons we obtained by the primers used. First, all OTU representative sequences were aligned against the 2 152 651 16S sequences in the SILVA 138.1 (Boratyn et al., 2013). The SILVA sequences have a median length of 1 403 bases and range in length from 900 to 4 000 bases. OTUs from subset A achieving a best match with ≥ 95% identity over ≥ 95% of their length were marked as ‘known’, i.e. resembling something already known from the SILVA database. The full-length corresponding matching SILVA sequences were then retrieved as representative for‘ known ’OTUs (Fig. 1, box Q1).

It should be noted that many of the seed sequences used for reconstruction (see Section: Reconstructing 16S using METASEED) derive from the same‘ known ’OTUs, obfuscating the need for sequence reconstruction as matching SILVA sequences are already known. The reconstruction was, however, still performed in order to test how closely the reconstructed sequence could be matched to the corresponding SILVA sequence.

### Reconstructing 16S using METASEED

OTUs were used as ‘seed sequences’ within the METASEED workflow to reconstruct a longer section of the 16S rRNA gene from shotgun metagenome reads. METASEED collects reads from the shotgun metagenome data, aligns them to the seed sequences and combines assignment results with abundance information. For each seed, a reconstructed sequence is reported if the sequence is ≥ 500 bp in length and the alignment occurs with ≥99% identity to the corresponding seed. Detailed information on our METASEED approach can be referred from the recent publication (Philip et al., 2024).

Reconstructed sequences were subsequently aligned to the SILVA database, using the same thresholds as above. As mentioned before (see Section: SILVA alignments), the ‘known ’ seeds could now be checked to verify whether the seed and its reconstructed sequence matched the same SILVA sequence, or at least the same taxon. The most interesting reconstructed sequences derive from cases where the SILVA database generated no matches to the seed nor to its reconstructed sequence. These reconstructed sequences were then added to our collection as new 16S sequences, representing new taxa relative to the SILVA collection (Fig. 1, boxes Q2 and Q3).

### Shotgun data assembly

The shotgun metagenome data from the 94 samples in subset A were filtered and merged using BBduk and BBmerge (Bushnell, 2014) from BBmap v39.01. BBDuk parameters included k-mer size 23 (k=23), hamming distance 1 (h_dist=1), 3’ end trimming quality 20 (Q_trim=20), and read discarding for average quality below 20 (Q_discard=20). No trimming was applied to the 5’ end (trim_left=0), bases beyond position 150 were trimmed (trim_right=150), and reads shorter than 30 bases were discarded (min_len=30). The assembly was done using metaspades v3.15.5, executed within a Singularity container (Nurk et al., 2017). Inputs included paired-end, merged, and unpaired reads, with assembly parameters configured for memory allocation, thread usage, and temporary file handling. The `--only-assembler` option was used to bypass read error correction.From the assembled contigs we used Barrnap v0.9 (https://github.com/tseemann/barrnap) to extract 16S sequences from the contigs using length threshold ≥ 800 bases. As 16S sequences extracted from different samples may be identical, all 16S sequences were clustered using VSEARCH v2.22.1, as described below, to obtain a resolution of 95% identity (Fig. 1, box Q4).

Finally, we also included 16S centroids that didn’t give any match to the above sequences. These unmatched 16S sequences represent the lowest quality part of the reference database (Fig. 1, box Q5).

### Taxonomic assignments

Taxonomic assignments of all database sequences was obtained using the SINTAX algorithm (Edgar, 2016) implemented in the VSEARCH software, where we used the GSR database and the NCBI taxonomy (Molano et al., 2024). This means sequences were assigned to all ranks to species, with confidence scores at each rank.

### Phylogenetic placement of METASEEED and Barrnap derived sequences

Phylogenetic placement of the METASEED-and Barrnap-derived sequences were determined since these are ‘new’ 16S variants. First, all database sequences (excluding short amplicons) were aligned and filtered using `cmsearch` in the `infernal` software v1.1.4 (Eddy, 2002) and secondary structure model for bacterial 16S downloaded from the `Rfam` database (Kalvari et al., 2021). This level of filtration, utilizing `cmsearch`, was applied to eliminate potential low-quality sequences from the database and short amplicons were excluded to minimize potential noise in this analysis. Sequences with an overall score (bitscore/length of the sequence) below 0.9 were filtered out. A multiple alignment of the resulting sequences was created using `cmalign` and computed evolutionary distances using Felsenstein-81 model (Felsenstein, 1981) and the `ape` R package (Paradis & Schliep, 2019). These distances were then used in a hierarchical clustering with average linkage and the sequences were split into 64 clusters. The clusters with at least 5 members were subject to a Fisher exact test for enrichment of sequences originating from either METASEED or Barrnap, i.e. sequences not previously found in SILVA. We considered a cluster to be enriched with new variants, if the odds-ratio is greater than 1 and the p-value is less than 0.05. In these clusters there are some 16S variants where the taxonomy is known to phylum rank, and the dominating phyla in the enriched clusters are mentioned in the results.

### Database comparisons

As previously stated, one of the main motivations behind this work is to improve the database that will be used to analyses amplicon data from Oxford Nanopore Technology. EMU (Curry et al., 2022) is a standard software for profiling such data and we wanted to compare our database to the default EMU database. We first used VSEARCH alignments with default settings to assess the overlap between our generated database and the default EMU database. Matches that failed to show at least 95% identity were not considered.

Next, we wanted to test how the Nanopore reads from the 90 subset B sediment samples would be profiled using the EMU software in combination with its default database (EMU-DB) as well as our custom database (AQUAeD-DB) using default settings. In addition, we also tested the profiling of Illumina de-replicated reads using standard VSEARCH alignments, i.e. pairwise alignment between read and database sequences, and assigning to the best, an identity threshold of 95% was used. The EMU output was processed using an R script to convert it into the read count matrix format. For VSEARCH, the read count matrix is provided if specified in the command. The read count matrices from the Illumina VSEARCH and Nanopore EMU tool assignments for the AQUAeD-DB and EMU-DB databases were used to compute relative abundance. Prior to the correlation analysis, missing database sequences from the Illumina read count were added to the Nanopore read count matrix, and vice versa, with a count of zero for each sample. This was done to resolve disparities in matrix lengths.

## Results

### Addition of SILVA sequences to the database

The full-length sequences were first collected from the SILVA database, by matching them against the16S rRNA amplicon representative sequences, generated by conventional short-read data processing. Using a 95% identity threshold, our 16S rRNA pipeline generated 14,393 OTUs from the subset A data. Each OTU has a representative sequence consisting of 400 - 440 bases in accord with the length of the V3-V4 amplicons. These sequences were then aligned against the SILVA database using VSEARCH. In total 5 278 of OTUs showed at least 95% identity matches with some of the SILVA sequences (identity distribution shown in Fig. 2). This represents 84% - 96% of the total OTU abundance in these 94 samples. The 4 921 distinct SILVA sequences matched to the OTU sequences were collected and filtered using `infernal cmsearch,` which resulted in 4 838 sequences. The SILVA sequences were added to the database as full-length representatives of these OTUs and they belong to the highest quality (Table 1, Q1) sequences in the database.

**Fig. 2.**
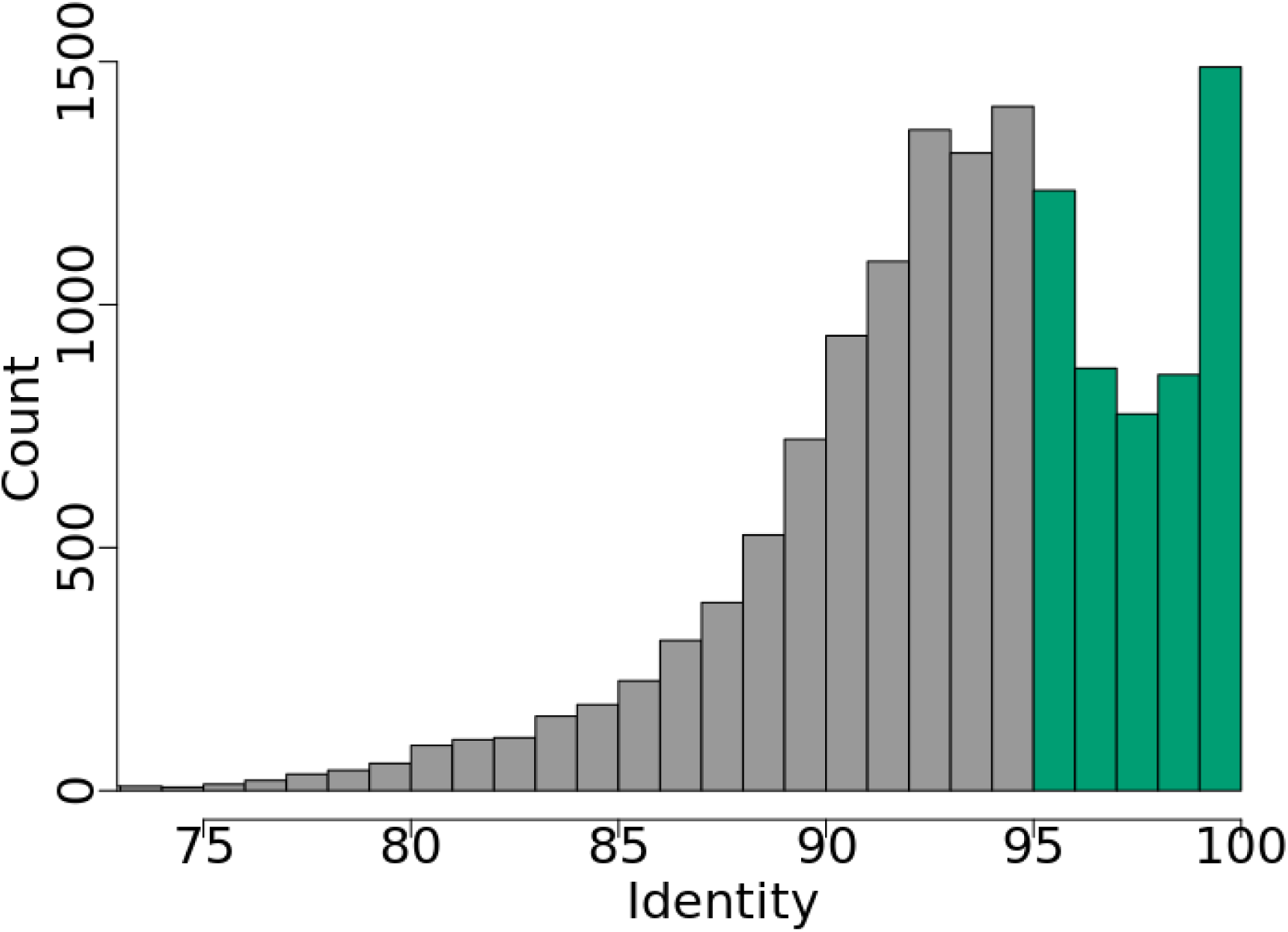
Quality of OTU identifications. Frequency distribution of OTU sequences (y-axis) with varying degrees of sequence similarity (x-axis) to SILVA database sequences. The green columns indicate the number of OTUs with a SILVA match while the grey columns indicate no relevant match to SILVA database.

**Table 1:** The AQUAeD-DB contains sequences of 5 quality categories, from the best (Q1) to the poorest (Q5) Category descriptions are provided in Materials and Methods. Lengths are shown in basepairs (bp).

| Category | Origin | Number of sequences | Minimum length | Median length | Maximum length |
| --- | --- | --- | --- | --- | --- |
| **Q1** | SILVA | 4838 | 900 | 1461 | 1668 |
| **Q2** | High Quality METASEED | 311 | 800 | 918 | 1522 |
| **Q3** | Low Quality METASEED | 406 | 501 | 688.5 | 799 |
| **Q4** | 16S from contigs | 320 | 800 | 924 | 1577 |
| **Q5** | **Amplicons** | **8670** | **288** | **426** | **438** |

### Reconstructing 16S rRNA genes using METASEED

All 14 393 OTUs were passed to the METASEED together with shotgun metagenome data in order to re-construct near full-length 16S rRNA from the shorter amplicon sequences. The screening process for the METASEED results is elaborated in Fig. S1. This analysis resulted in 3 525 16S rRNA sequences, of which 3 392 were ≥ 500 bases in length. Of these, 2 947 had at least 99% sequence identity to their seeds, and were considered re-constructed 16S sequences.

From the 2 947 reconstructed 16S sequences, 1 176 had no match to SILVA sequences already collected (≥ 95% identity). Reconstructed 16S rRNA sequences with a minimum length of 800 bases were regarded as high quality, while sequences 500 - 800 bases in length were considered low quality. This resulted in 421 high quality and 755 low quality reconstructed sequences. The total of 1 176 METASEED reconstructed sequences were filtered using `cmsearch`, which led to the addition of 717 reconstructed sequences to the database. The Q2 and Q3 category in Table 1 summarize these results. Fig.S2 shows the increase in captured relative abundance when Q1+Q2+Q3 sequences are included, compared to using only Q1.

### Addition of 16S rRNA from contigs using Barrnap

We also looked for 16S rRNA genes in the assembled contigs from the metagenome data using the `barrnap` software. From the 94 samples, we found in total 9 839 short or longer sequences recognized as 16S rRNA. The number of sequences was reduced to 1 149 when applying a length threshold of at least 800 bp. Finally, since the same 16S rRNA sequence may have been assembled and extracted from different samples, these sequences were clustered using a 95% identity threshold, which reduced the number of sequences to 518. Again, we used the `cmsearch` filtering, resulting in 320 high quality near full-length 16S rRNA sequences, which were added to database. The total length of assembled contigs was greater than 32 000 megabasepairs, implying that roughly one full-length high-quality 16S rRNA was found for every 100 mega base-pairs of genomic sequence (Table 1, Q4).

Apart from the sequences collected from the above-mentioned sources, we also included OTU sequences that did not match to any sequences added to the database from SILVA, METASEED, and Barrnap when using a 95% identity threshold. This 8 670 OTUs accounts for the Q5 category(Table 1). Including all categories (Table 1), the final size of the AQUAeD database was 14 545 16S sequences.

### Taxonomic assignments

All sequences in the AQUAeD database were classified using `SINTAX` (Fig. 3), with the exception of sequences belonging to the Q1 category, which were collected from the SILVA database with taxonomic assignments.

**Fig. 3.**
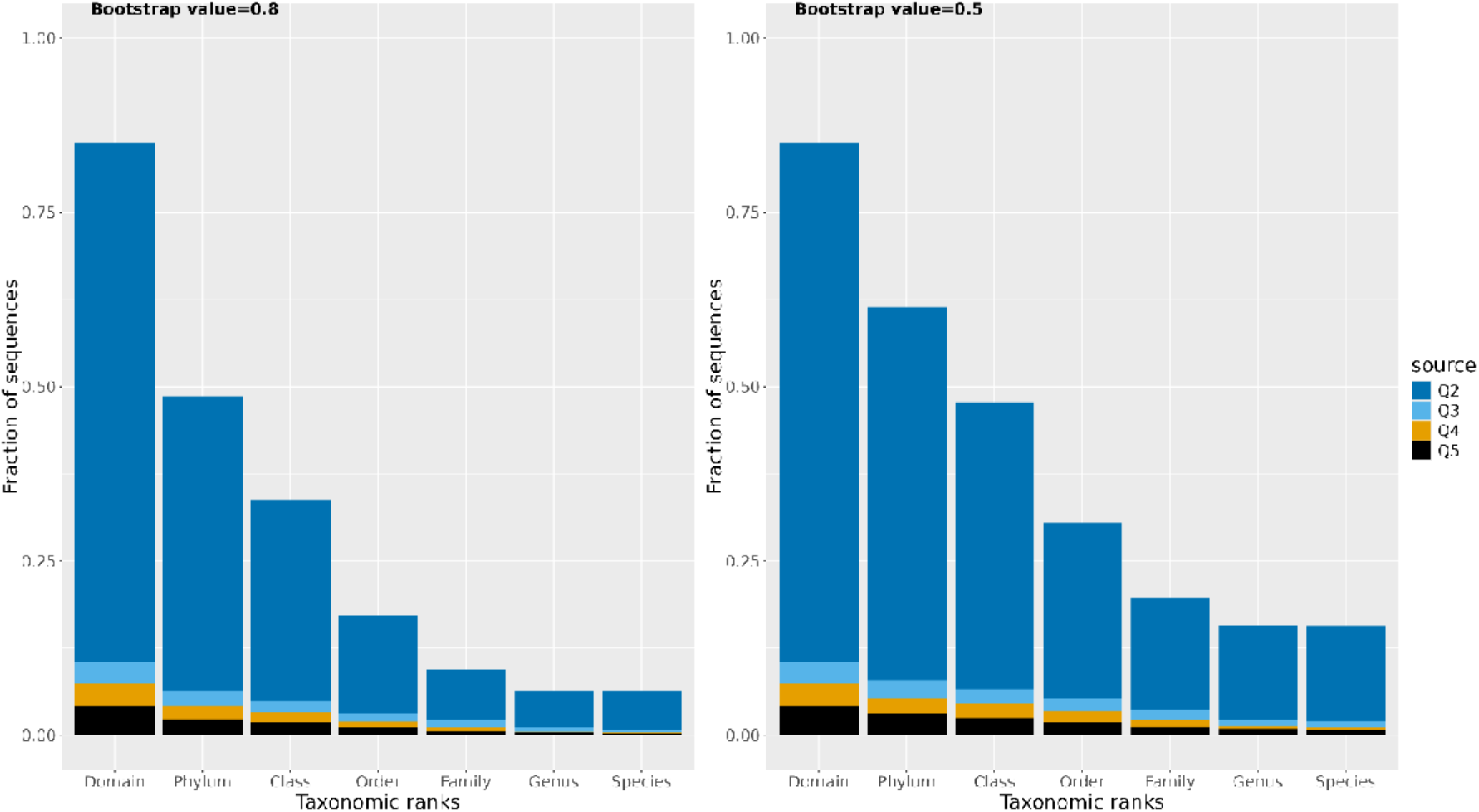
The bars depict the fraction of classified sequences (Q2 - Q5) in the AQUAeD Database using SINTAX (bootstrap value 0.8 and 0.5) and the GSR databases. The x-axis shows various taxonomic ranks, and y-axis shows the fraction of classified sequences.

### Enrichment of ’new ’16S variants

Sequences in categories Q2, Q3 and Q4 (Table 1) are ‘new’ 16S rRNA gene variants in the sense that they are not included in the SILVA database. We wanted to see how these distribute when clustered among the sequences in Q1 as well. The multiple sequence alignment of these 5 875 sequences was followed by the evolutionary distance calculation, cluster generation and using the Fisher test for enrichment of the ‘new variants’. From 64 clusters we found that 17 clusters were enriched by ‘new’ variants. The other members of these enriched clusters predominantly consisted of phyla Planctomycetota, Myxococcota, and Verrucomicrobiota (Table 2).

**Table 2:** Enrichment of ‘new’ 16S variants. Results of Fisher tests for 17 ‘new variants ’enriched clusters when k=64.

| **Odds ratio** | **Cluster size** | **New variants fraction** | **Dominant phylum** |
| --- | --- | --- | --- |
| 7.06 | 20 | 0.6 | Planctomycetota |
| 7.01 | 5 | 0.6 | Desulfobacterota,Proteobacteria |
| 6.56 | 12 | 0.58 | Patescibacteria |
| 4.015 | 13 | 0.46 | Patescibacteria |
| 3.97 | 35 | 0.46 | Spirochaetota |
| 3.27 | 39 | 0.41 | Verrucomicrobiota |
| 3.01 | 23 | 0.39 | Spirochaetota |
| 2.71 | 30 | 0.37 | Verrucomicrobiota |
| 2.59 | 62 | 0.35 | Chloroflexi |
| 2.58 | 114 | 0.35 | Verrucomicrobiota |
| 2.55 | 153 | 0.35 | Planctomycetota |
| 2.54 | 37 | 0.35 | Firmicutes |
| 2.34 | 30 | 0.33 | Firmicutes |
| 2.34 | 27 | 0.33 | Fibrobacterota |
| 2.13 | 61 | 0.31 | Planctomycetota |
| 2.07 | 72 | 0.31 | Verrucomicrobiota |
| 1.61 | 241 | 0.25 | Myxococcota |

### Database comparison

EMU software is the standard tool used for taxonomic profiling of Nanopore 16S reads. It comes with a default database (EMU-DB), consisting of 49 243 sequences. From a comparison between the EMU-DB and AQUAeD-DB, we observed that 18 584 (38%) EMU database sequences had a match to 509 (3.5 %) AQUAeD database sequences when using a 95% identity threshold.

The 88 subset B samples resulted in a total of 973 758 dereplicated Illumina reads, corresponding to 51 397 distinct sequences. The alignment of these reads (95% identity) to the EMU-DB resulted in a match for 3% of the database sequences, comprising 52% of the total Illumina reads. Alignment with the AQUAeD-DB yielded a match to 23% of the database sequences, covering 99% of the total Illumina reads. The Nanopore amplicon reads from the 88 subset B samples having length between 800 and 1 400 bases resulted in 4 972 056 reads. The analysis of Nanopore reads resulted in 3 967 AQUAeD database sequences, accounting for approximately 27% of the database, while the Nanopore reads matched to 2560 EMU-DB sequences, accounting for approximately 5% of the database.

A comparison of read count matrices revealed that 82% of the EMU-DB sequences in the Illumina read count matrix were missing in the corresponding Nanopore read count matrix, while 17% of the sequences in the Nanopore matrix were the missing from Illumina matrix. For the AQUAeD-DB read count matrices, the overlap was notably better, with 45% of Illumina sequences and 13% of Nanopore sequences missing from their counterparts, indicating a lower level of missing database sequences. Correlation analysis demonstrated high positive correlation between read count matrices for the AQUAeD-DB, whereas the EMU-DB shows very weak negative correlation (Fig. 5). The relative mean abundance of Illumina and Nanopore results for AQUAeD-DB demonstrate a significant positive correlation (Fig. 6).

## Discussion

The motivation for building the AQUAeD-DB was to enhance the use of Nanopore data in the study of seafloor sediments. Thus, we wanted to see if the use of AQUAeD-DB has some benefit compared to a standard database. Since the EMU tool is a popular choice for Nanopore 16S data processing, we used its default database (EMU-DB) in our comparison. In addition to using EMU for assigning the reads, we also used VSEARCH, which is typically used for Illumina data, just for comparison. The VSEARCH alignment directly between the AQUAeD-DB and EMU-DB reveals that 3.5% of the AQUAeD database could represent 38% of the EMU database, implying that the AQUAeD-DB, despite being smaller in the number of sequences, AQUAeD-DB covers an order of magnitude more seafloor diversity compared to EMU-DB. This difference could also be due to the EMU-DB containing more sequence variants from the same taxa, specifically from human-associated environments, which may be less relevant to seafloor ecosystems. This underscores the importance of utilizing environment-specific databases in monitoring studies.

The AQUAeD-DB comprises of 14 545 16S sequences that were collected from seafloor sediments ( Fig. 1). The Illumina amplicon sequencing of subset A seafloor samples yielded 14 393 centroid sequences when clustered at the coarse level of 95% identity. Approximately 40% of these sequences matched the SILVA database sequences, even with relatively liberal identity threshold of 95% ( Fig. 2). However, this set of matching SILVA sequences, the Q1, makes up the backbone of our AQUAeD-DB. To add more longer sequences to the database, we used our METASEED tool (Philip et al., 2024) to extend the amplicon sequences, making use of the Illumina metagenome data from the same samples (subset A). This resulted in the Q2 and Q3 part of the database. The total abundance covered by the database sequences in each sample did not increase appreciably by adding these sequences, but the increase was larger for the exceptionally high diversity samples (high Shannon index of 5-7, Fig. S2). This typically allowed us to obtain 16S sequences for the low-abundance and ecologically significant organisms from phyla like Planctomycetota (table 2). In the Q4 part of the database, we tried to collect long 16S directly from metagenome assembled reads. The fact that we only collect one such sequence per 100 megabases of assembled contigs indicates how difficult it is to obtain such information from the assembly of short read metagenome data. The limitations of metagenome assembly, specifically the challenges in retrieving 16S sequences, have been previously addressed (Hiseni et al., 2022; Mise & Iwasaki, 2022; Shaiber & Eren, 2019).

The AQUAeD-DB can be roughly split into two parts: The ‘known ’sequences (Q1) and the ‘new variants ’(Q2-Q5, Fig 3). The fractions of taxonomic assignments were generally low, especially for the Q3-Q5 sequences. This emphasizes that new sequence variants in the AQUAeD database (Q2-Q5) have poor representation in a public database, like the GSR used here. Considering the poor taxonomic information, we wanted to see if phylogenetic distances between these new variants, and the Q1 sequences indicate a grouping of these new sequences, or if they are scattered around in the phylogeny. By clustering based on phylogenetic distances, we found several clusters where these new variants are over-represented, which include significant marine microorganisms from the phyla Planctomycetota, Myxococcota, and Verrucomicrobiota (Dell’Anno et al., 2021; Freitas et al., 2012; Wu et al., 2024)

A clear difference was evident when we did the assignment with both VSEARCH, which is normally used for Illumina data, and EMU, which is the standard choice for Nanopore (Fig 4). The VSEARCH tool aligned the high-precision Illumina reads, with EMU-DB assigning almost half of the reads and spreading them out among only 1 474 of the nearly 50 000 database sequences. For AQUAeD-DB, almost all reads are assigned to 3 301 database sequences. The Nanopore reads showed different results compared to the VSEARCH assignment of Illumina data, especially in terms of database coverage. One of the major contributing factors can be the significantly higher noise levels compared to Illumina data; this leads to reads being distributed across a much larger number of sequences in both databases (Magi et al., 2017; Santos et al., 2020). Another difference lies in the fact that the EMU tool uses Expectation-Maximization (EM) algorithm and results in non-integer read counts for each taxa/database sequence, while result in the assignment more than 100% of the reads as we sume the readcounts. This likely indicates an artifact of the fuzzy assignment process (Szoboszlay et al., 2023). From VSEARCH+Illumina and EMU+Nanopore, we see how more organisms are being given read counts in the AQUAeD-DB compared to the EMU-DB. This is the effect of AQUAeD-DB containing more relevant taxa for seafloor data, while EMU-DB is a generic collection and probably more relevant for use in medical research.

**Fig. 4.**
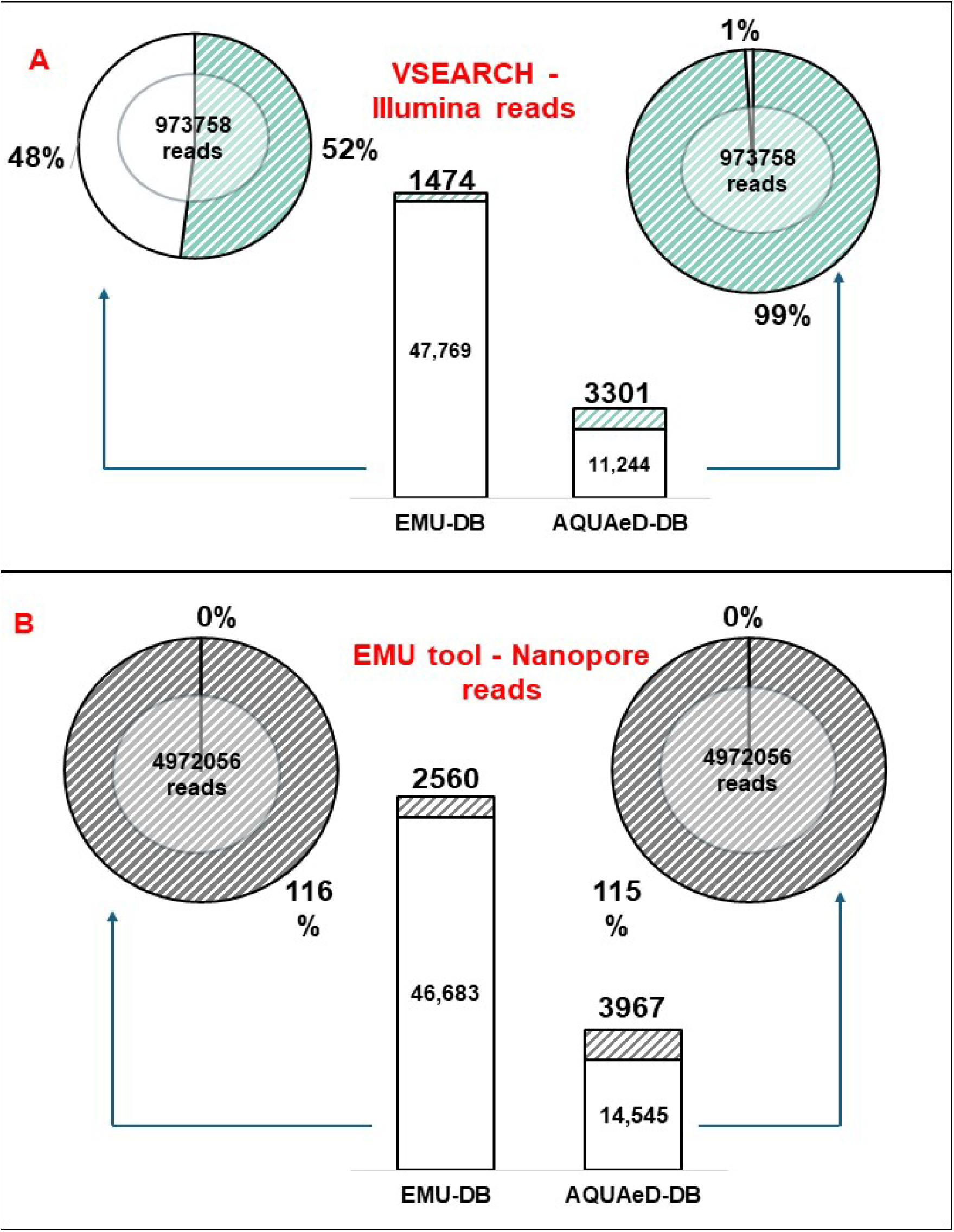
The Database comparison. A) Illumina results for taxonomic assignments using VSEARCH. B, Nanopore results for taxonomic assignments using EMU. B) the percentage reads assigned exceeds 100% due to the fuzzy assignment by EMU which resulted in a total ‘count ’of assigned reads that was larger than the number of reads. Pie charts represent reads and database sequences are represented by bars. Shaded/striped regions in each pie chart or bar indicate the fraction of reads assigned or the number of database sequences to which they were assigned, respectively. Detailed information about alignments can be found in Table S1.

The previously stated percentage of missing database sequences in the read count matrices for Illumina and Nanopore indicate that the EMU-DB analysis results exhibit greater disparities compared to the AQUAeD-DB results. The discrepancies here can be due to the different primers, possible primer bias, differences in the 16S rRNA regions used for Nanopore vs Illumina sequencing (Abellan-Schneyder et al., 2021; Wang et al., 2021; Zhang et al., 2024). However, clear positive correlation between Illumina and Nanopore relative abundance were revealed in AQUAeD-DB whereas EMU-DB shows very inconsistent and poor results (Fig 5). From these results it is evident that AQUAeD-DB results in more reliable and consistent assignments for both Illumina and ONT analysis. These findings were supported by the broad range of diversity, especially for the low-abundance taxa that the AQUAeD-DB database was able to cover (Fig 6). These observations once again underline the benefits of AQUAeD-DB and emphasize the potential implementation for Nanopore based environmental studies.

These findings indicated that AQUAeD-DB have illustrated a significant contribution in environmental analysis using ONT sequencing data. The database enrichment analysis implies that significant marine microbiota is represented in newly developed AQUAeD database, comprising both existing sequences and novel sequences from the aforementioned strategies. This optimizes taxonomical assignment and provides potential for more explicit environmental context. The study emphasizes the potential, and necessity of such databases in facilitating the implementation of the cutting-edge technologies, thus enabling the comprehensive environmental assessments.

**Fig. 5.**
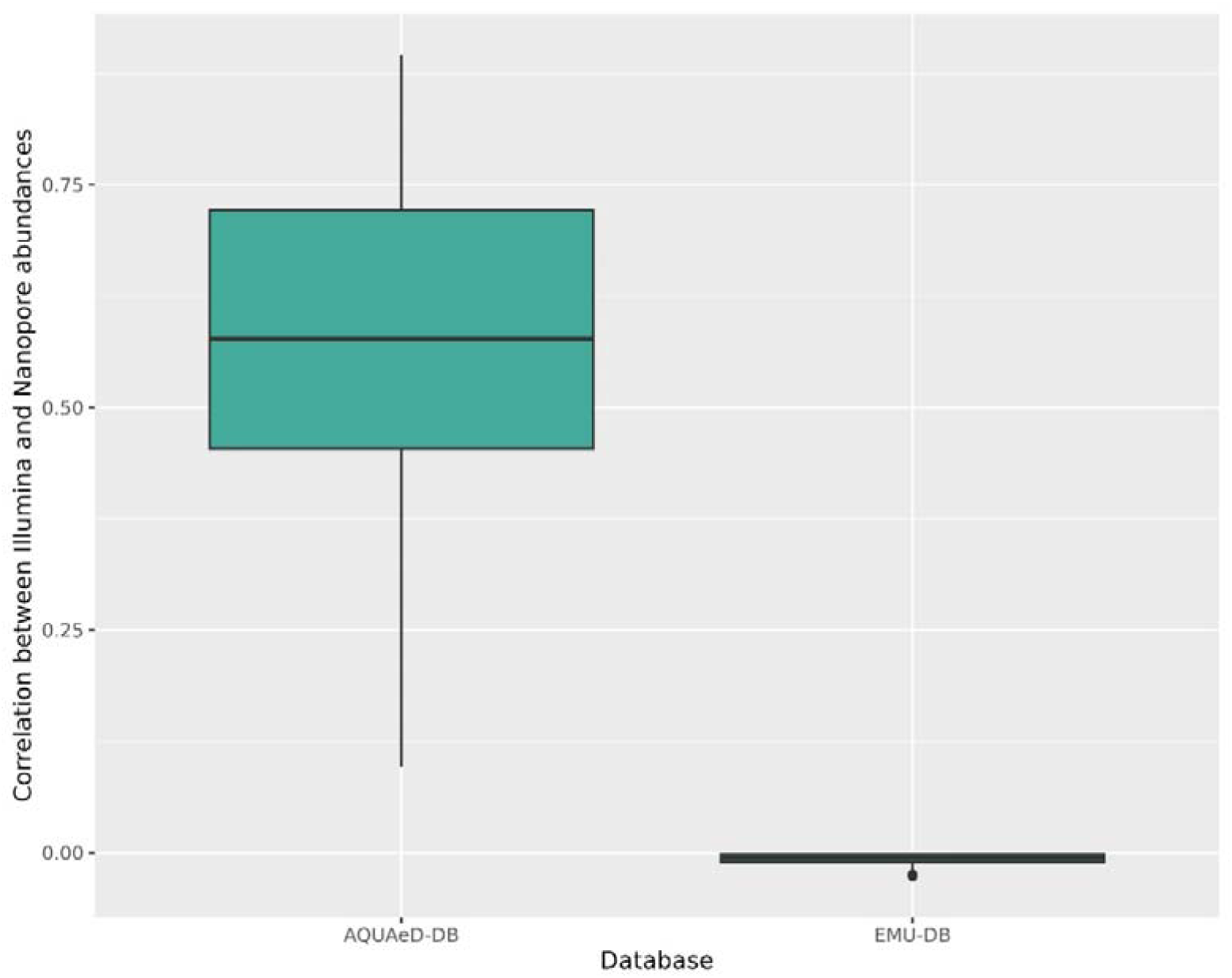
Correlation between Illumina and ONT read count matrices. Boxplots indicate correlation coefficients for read count matrix similarity when using the AQUAeD or EMU database.

**Fig. 6.**
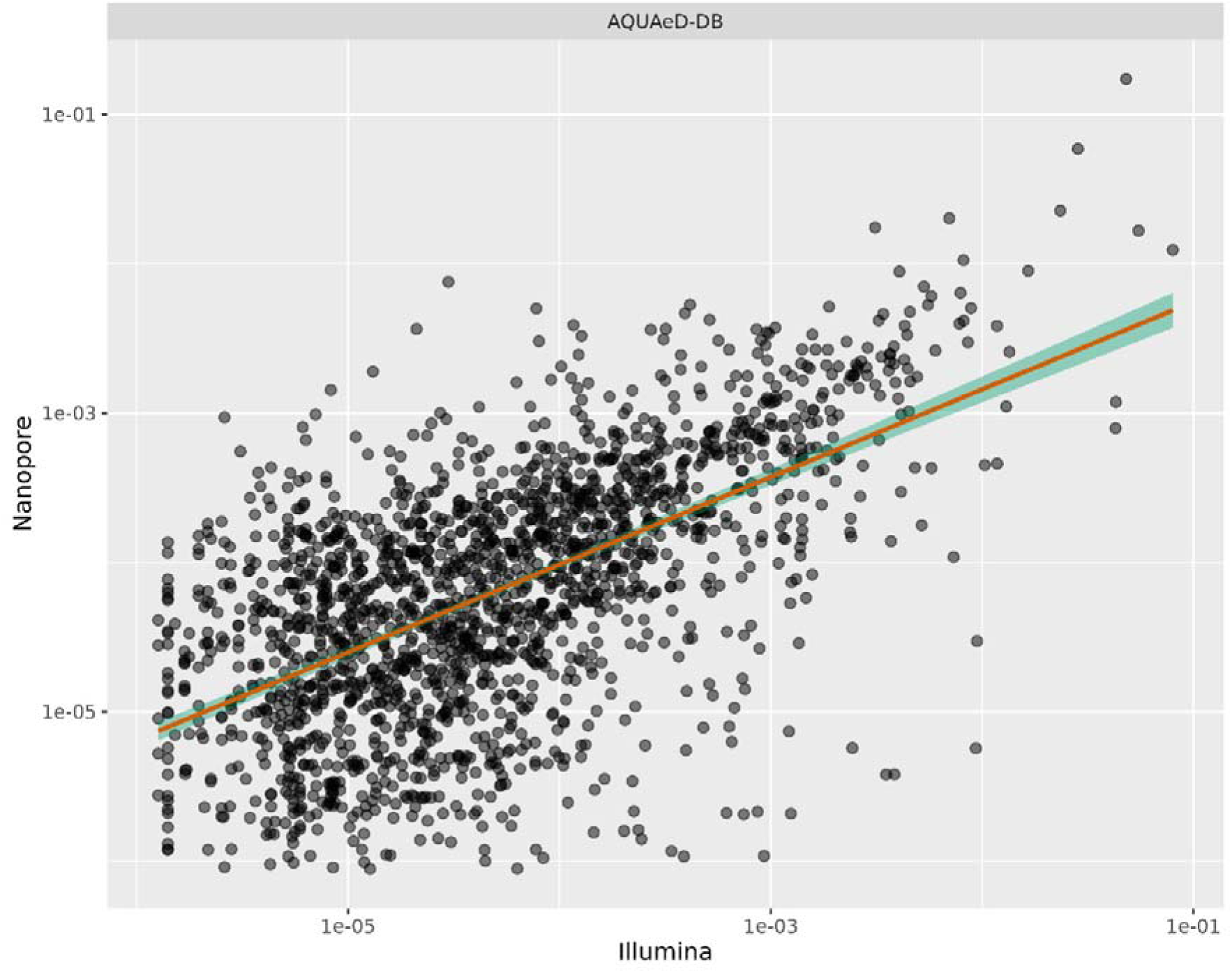
Relative mean abundance of Illumina versus Nanopore sequences for subset B samples. The scatterplot depicts the log of relative mean abundance of Nanopore (y-axis) versus the log of relative abundance of Illumina(x-axis) of AQUAeD-DB database sequences across samples. Cases where either Illumina or Nanopore values were zero are omitted.

## Conclusion

The newly developed AQUAeD-DB database is a specialized database for 16S rRNA gene sequences from highly diverse and complex environmental samples. The primary objective of AQUAeD-DB was to provide a better reference database for identification of Nanopore sequences targeting microbial communities in North Atlantic seafloor sediments. Based on results of the enrichment study and database comparison, this database is substantially diverse, and efficiently covers an expanded range of microorganisms that are poorly represented in other databases. This study was designed as a pilot project with the potential for future scalability of the database to incorporate additional 16S rRNA gene sequences. This expansion will be rooted in ongoing efforts to improve environmental surveys and monitoring using microbial profiles.

## Acknowledgements

We express our gratitude to the fish farmers that provided samples for this research. We would also like to thank the employees of STIM, Aqua Kompetanse, and Akvaplan-NIVA that performed the sampling.

## Data availability

The AQUAED-DB and taxonomy details are publicly available (https://arken.nmbu.no/~pmelcy/share/AQUAED-DB/). All raw sequences used in this study have been deposited in the NCBI BioProject database. Illumina 16S rRNA sequences from subset A and subset B is a part of the project with the accession code <u>PRJNA1128851</u>. The raw shotgun data of subset A has been uploaded with the accession code <u>PRJNA1128851</u>. The raw Nanopore sequencing data for subset B has been uploaded with the accession code <u>PRJNA1153974</u>. The metadata files for all datasets are available in https://arken.nmbu.no/~pmelcy/share/AQUAED-DB/metadata/

## Author contributions

Knut Rudi, Lars Snipen and Melcy Philip conceived the study; Jessica Louise Ray, Morten Stokkan, Ragnhild Pettersen and Sanna Majaneva design the collection of sediment samples; Tonje Nilsen performed all lab experiments; Lars Snipen and Melcy Philip performed all analyses, with conceptual input from Knut Rudi; Knut Rudi, Lars Snipen and Melcy Philip contributed to interpretation of results; Knut Rudi, Lars Snipen and Melcy Philip drafted the manuscript. All author(s) read and commented on the manuscript.

