## Supplementary files for "A Targeted Reference Database for Improved Analysis of Environmental 16S rRNA Oxford Nanopore Sequencing Data"

**Supplementary data**

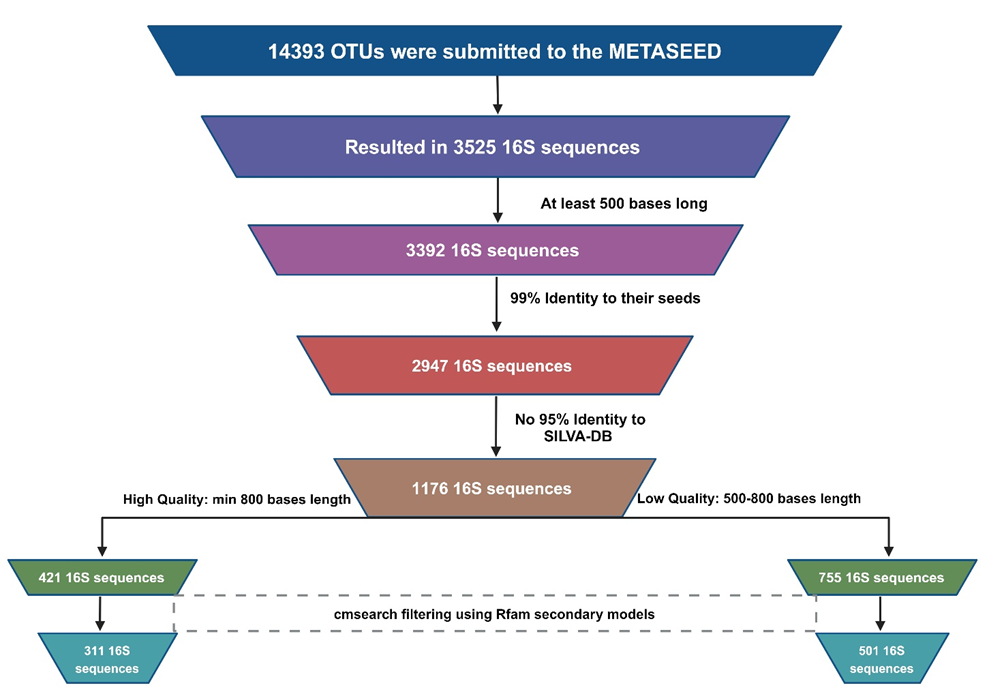

**Fig. S1** The Fig. depicts the workflow of the screening process in METASEED analysis

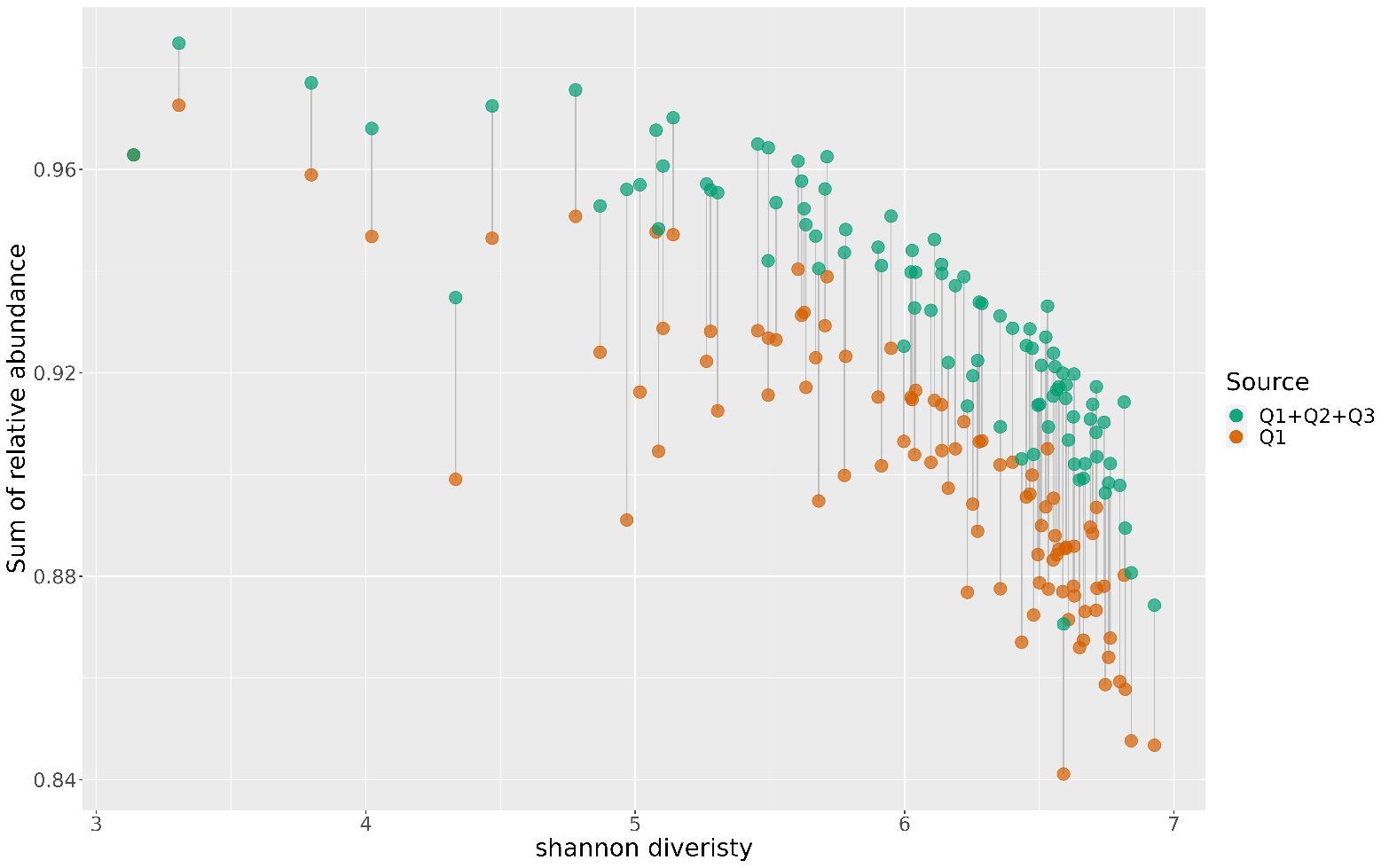

**Fig. S2** The Fig. depicts the sum of relative OTU abundance (y-axis) in each of the ninety-four samples covered by the SILVA set only (orange dots) and the increase obtained by adding the reconstructed sequences as well (green dots). The x-axis shows Shannon diversity for each sample. The distance between a brown and green dot shows the increase in relative abundance as we add the Q1 and Q2 sequences to the collection.

**Table S1:** The table presents the VSEARCH alignment results of Illumina dereplicated reads and EMU assignment results of Nanopore reads against EMU database and AQUAeD database.

|  | **Total** | **AQUAeD (14545)** | **EMU (49243)** |
| --- | --- | --- | --- |
| **Distinct Illumina Sequences** | 51397 | 3301 | 1474 |
| **Corresponding Illumina Reads** | 973758 | 966477 | 358760 |
| **Illumina Reads Percent** | 100 | 99 | 52 |
| **Number of total reads and unique database sequence matches (Nanopore)** | 49 72 056 | 3967 | 2560 |
| **Fraction of database sequences (Nanopore)** | 100 | 27 | 5 |
